# Stochastic Sampling of Structural Contexts Improves the Scalability and Accuracy of RNA 3D Module Identification

**DOI:** 10.1101/834762

**Authors:** Roman Sarrazin-Gendron, Hua-Ting Yao, Vladimir Reinharz, Carlos G. Oliver, Yann Ponty, Jérôme Waldispühl

## Abstract

RNA structures possess multiple levels of structural organization. Secondary structures are made of canonical (i.e. Watson-Crick and Wobble) helices, connected by loops whose local conformations are critical determinants of global 3D architectures. Such local 3D structures consist of conserved sets of non-canonical base pairs, called RNA modules. Their prediction from sequence data is thus a milestone toward 3D structure modelling. Unfortunately, the computational efficiency and scope of the current 3D module identification methods are too limited yet to benefit from all the knowledge accumulated in modules databases. Here, we introduce BayesPairing 2, a new sequence search algorithm leveraging secondary structure tree decomposition which allows to reduce the computational complexity and improve predictions on new sequences. We benchmarked our methods on 75 modules and 6380 RNA sequences, and report accuracies that are comparable to the state of the art, with considerable running time improvements. When identifying 200 modules on a single sequence, BayesPairing 2 is over 100 times faster than its previous version, opening new doors for genome-wide applications.

## 1 Introduction

RNAs use complex and well organized folding processes to support their many non-coding functions. The broad conservation of structures across species highlights the importance of this mechanism [35,14]. RNAs can operate using folding dynamics [25] or hybridization motifs [2]. Yet, many highly specific interactions need sophisticated three dimensional patterns to occur [15,11,13].

RNAs fold hierarchically [36]. First, Watson-Crick and Wobble base pairs are rapidly assembled into a secondary structure that determine the topology the RNA. Then, unpaired nucleotides form non-canonical base pairs interactions [16], stabilizing the loops while shaping the tertiary structure of the molecule. These non-canonical base pairing networks have thus been identified as critical components of the RNA architecture [4] and several catalogs of recurrent networks along with their characteristic 3D geometries are now available [10,7,27,28,30,12]. They act has structural organizers and ligand-binding centers [8] and we call them *RNA 3D modules*.

In contrast to well-established secondary structure prediction tools [20,22], we are still lacking efficient computational methods to leverage the information accumulated in the module databases. Software such as RMDetect [8], JAR3D [34] and our previous contribution BayesPairing 1 [32] have been released, but their precision and scalability remains a major bottleneck.

The significance of a module occurrence is typically assessed from recurrence: substructures that are found in distinct RNA structures are assumed to be functionally significant [30]. Based on this hypothesis, three approaches have been developed so far for the retrieval and scoring of 3D modules from sequence. The first one, RMDetect, takes advantage of Bayesian Networks to represent base pairing tendencies learned from sequence alignments. Candidate modules found in an input sequence are then scored with Bayesian probabilities. However, while showing excellent accuracy, RMDetect suffers from high computational costs, and minimal structure diversity among modules predicted [32]. Another option is JAR3D [34], which refined the graphical model-based scoring approach introduced by RMDetect and represents the state of the art for module scoring. However, it was not designed to maximize input sequence scanning efficiency and is limited in module diversity, only being applied to hairpin and internal loops. Finally, BayesPairing 1 [32], a recently introduced tool combining the Bayesian scoring of RMDetect to a regular expression based sequence parsing, is able to identify junction modules in input sequences and showed improved computational costs compared to RMDetect, which it was inspired from. Unfortunately, none of the aforementioned software can be used for the discovery of many RNA 3D modules in new sequences at the genome scale.

In this paper, we present BayesPairing 2, an efficient tool for high-throughput search of RNA modules in sequences. BayesPairing 2 analyzes the structural landscape of an input RNA sequence through secondary structure stochastic sampling and uses this information to identify candidate module insertion sites and select modules occurring in a favorable structural context. This pre-scoring stage enables us to dramatically reduce the number of putative matches and thus to (i) simultaneously search for multiple modules at once and (ii) eliminate false positives. BayesPairing 2 shows comparable performance to the state of the art while scaling gracefully with the number of modules searched. It also supports alignment search, a feature of RMDetect which could not be integrated in the BayesPairing 1 framework. All these improvements support potential applications at the genome scale.

## 2 Methods

### Concepts and model

A **non-canonical 3D module** consists in a set of non-canonical base pairs [17]. Modules occur within a **secondary structure loop**, consisting of one or several stretches of unpaired positions within an RNA transcript, also called **regions**, delimited by classic Watson-Crick/Wobble base pairs.

At the **thermodynamic equilibrium**, an RNA sequence *w* is expected to behave stochastically and adopt any of its secondary structure *S*, **compatible** with *w* with respect to canonical Watson-Crick/Wobble base pairing rules, with probability proportional to its **Boltzmann factor**[23]. The **Boltzmann probability** of a secondary structure *S* for an RNA sequence *w* is then

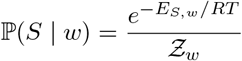

where *E_S,w_* represents the free-energy assigned to the (*S, w*) pair by the experimentally established Turner energy model [37], 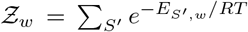 is the partition function [23], *R* is the Boltzmann constant and *T* the absolute temperature. By extension, the Boltzmann probability of a given loop to occur within a sequence *w* is simply defined as

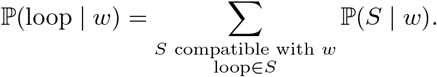

In the current absence of thermodynamic data for non-canonical base-pairs and modules, we adopt a probabilistic approach, and model the sequence preferences associated with a module statistically as a **Bayesian network**, following Cruz *et al* [8]. The structures of Bayesian networks are systematically derived from the base pairs occurring within **recurrent 3D motifs**[30]. Such motifs are typically mined within available 3D RNA structures in the PDB [5], and clustered geometrically.

Networks are then decomposed in a way that minimizes direct dependencies between individual positions of the module, while transitively preserving the emission probabilities. As illustrated in Figure 2, we use a **tree decomposition**[6] of the network to minimize the maximum number of prior observations at each position, a strategy shared by instances of the junction tree methods [3]. Maximum likelihood conditional emission probabilities are then learned for each module using pseudo-counts.

**Fig. 1:**
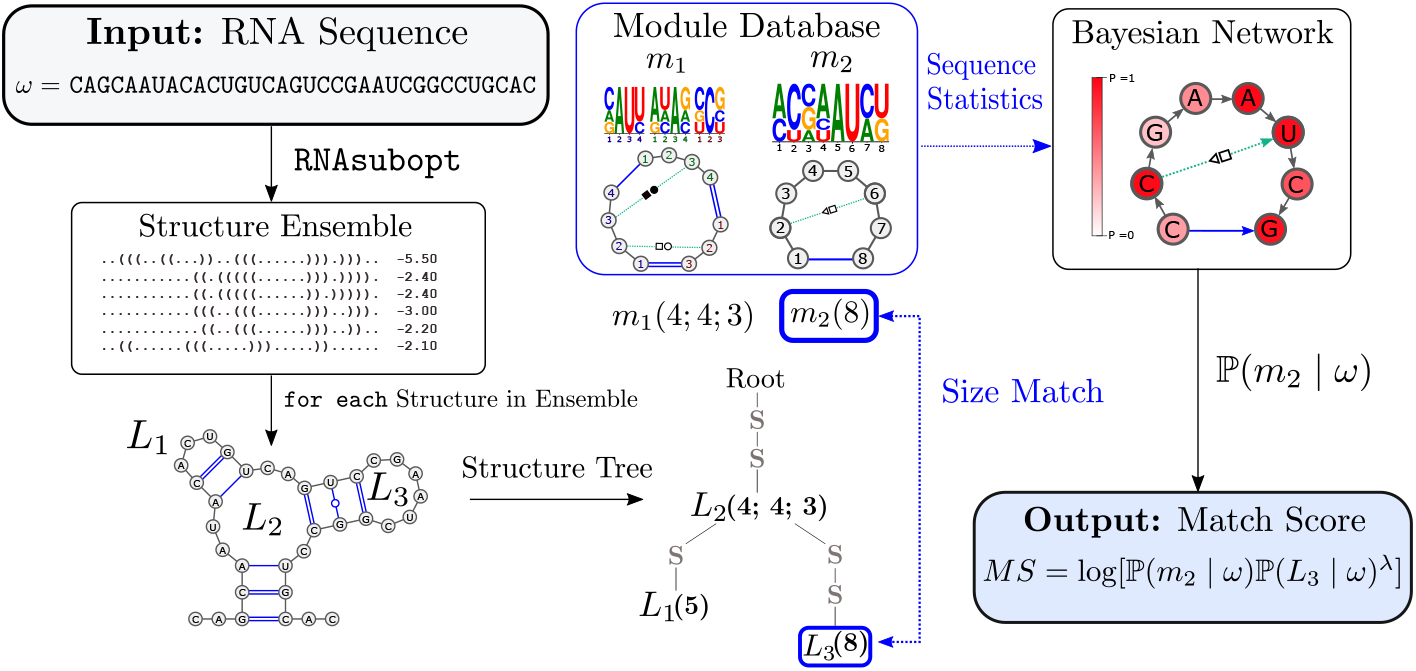
The BayesPairing 2 workflow addresses the identification of noncanonical 3D modules, *i.e.* arrangements of canonical and non canonical base pairs that are essential to the 3D architecture of RNAs. It takes as input either an RNA transcript or a multiple sequence alignment, possibly supplemented with a (shared) secondary structure, and returns an ordered list of occurrences for candidate modules. Its key idea is to match predicted secondary structure loops, highly likely to occur in thermodynamically-stable models, against a database of local modules learned from sequence data filtered for isostericity [19]. In this figure, we show the identification pipeline for one module on one structure of the ensemble. This is then repeated for all modules, for all structures.

**Fig. 2:**
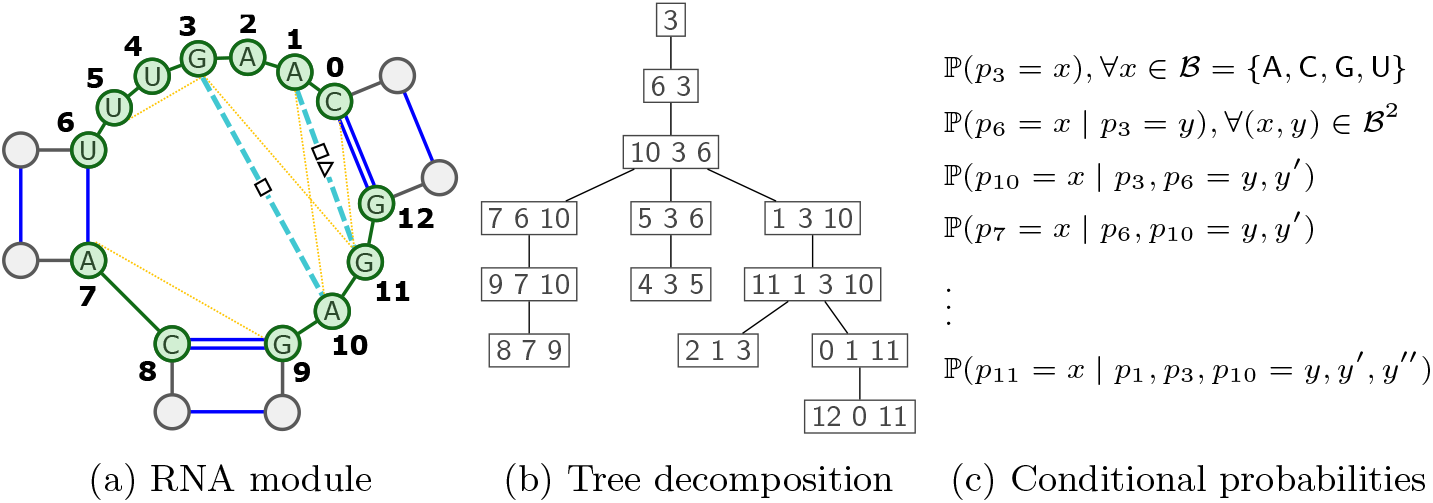
An RNA 3D module (2a), here the three-way junction of the TPP riboswitch, represented in green, drawn in its structural context. Dashed and dotted lines respectively represent non-canonical base pairs and stacking interactions. A tree decomposition (2b) of the module represents the dependencies between the module positions, leading to conditional probabilities (2c), estimated from available sequence data

The **emission probability** for the positions of a module *m* to be assigned to a nucleotide content *A* is then given by

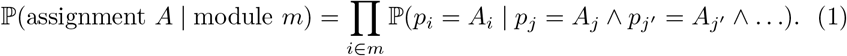

where *p_j_*, *p_j_′,…* represent the content of positions *j, j′,…*, the positions conditioning the content *p_i_* of position *i*, as derived using the tree decomposition, and *A_i_* represents the content of the *i*-th position in *A*. Using Bayes Theorem while assuming uniform priors for both assignments and modules 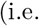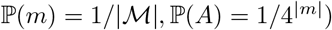, we obtain

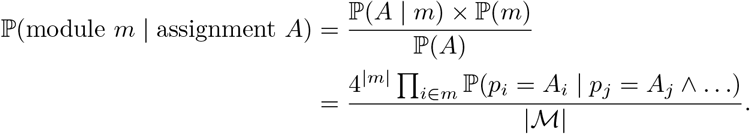

where 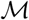 represents the set of admissible modules.

The final **match log-odds score** MS associated with a motif *m* being embedded within a given loop (*i.e.* at a given position) for an RNA sequence *w* is given by

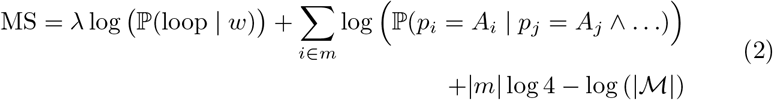

where λ is a term that allows to control the weight of the structure and local sequence composition.

### Algorithmic considerations and complexity

On an algorithmic level, for given sequence *w* and module *m*, we remark that it suffices to optimize for the first two terms of the above equations, the others being constant for a given module. A list of loops having highest Boltzmann probability 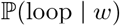 is first estimated from a statistical sample, generated using (non-redundant) stochastic backtrack [9,24,31]. The second term, *i.e.* the probability of the module content, is only evaluated for the loops that are compatible with the size constraints of the module, with tolerance for a size mismatch of up to one base per strand (—∞ otherwise). Its evaluation uses conditional probabilities, learned from a tree-decomposition of the module, as described in Figure 2. Matches featuring scores higher than a **cut-off** *α* are then reported as candidates.

The **overall complexity** of the method, when invoked with a module *m* and a transcript *w* of length *n* is in 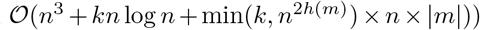, where *k* denotes the number of sampled secondary structures and *h*(*m*) is the total number of helices in *m*. It follows a sequence-agnostic precomputation in 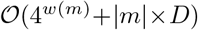, where *w*(*m*) represents the tree-width of *m*, and *D* represents the overall size of the dataset used for training the model.

Remark that, while our reliance on sampling formally makes our method a heuristic in the context of optimizing the objective in Equation (2), it must be noted that sampling provides a **statistically consistent** estimator for the probabilities of loops. Moreover, the probabilities associated with all possible loops could be computed exactly using constrained dynamic programming in time 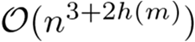 [20].

### Implementation

Secondary structures are non-redundantly sampled from the whole ensemble if the structure is not provided in the input, using RNAsubopt for a single sequence, or RNAalifold for a set of pre-aligned sequences [20,24,31]. Tree decompositions of modules are computed by the htd library [21] and conditional probabilities are learned using pgmpy [1]. BayesPairing 2 is freely available as a downloadable software at (http://csb.cs.mcgill.ca/BP2).

### Positioning against prior work

Using stochastic sampling in BayesPairing 2 allows to efficiently score all modules of a dataset in a single sequence search, unlike the previous version, which requires multiple regex searches on the sequence for each module. While searching structure-first improves the sensitivity, especially on modules without a strong sequence signal, it can add potential false positives, especially for small modules which appear a lot in secondary structures. This translates into more candidates scored, but scoring a candidate is much faster than scanning a sequence. Thus, BayesPairing 2 is much more more efficient when searching for many modules. In addition, the ability to sample with RNAalifold allows BayesPairing 2 to take full advantage of aligned sequences.

## 3 Results

### 3.1 Rna3Dmotif dataset

In order to assess the performance of BayesPairing 2 on its own and in context with that of BayesPairing 1, we assembled a representative sequence-based dataset of local RNA 3D modules. We ran Rna3Dmotif on the non-redundant RNA PDB structure database [18]. Identified modules were then matched to Rfam family alignments via 3D structure positions. Sequences from these alignments were filtered to remove poorly aligned sequences, using isostericity substitution cutoffs ensuring that the extracted sequences could adopt their hypothesized structure. Modules matched to at least 35 sequences were added to the dataset. 75 modules, totaling 20 125 training sequences, were collected. To assess the presence and potential impact of **false positives (FP)** and **true negatives (TN)**, a negative dataset was assembled. To build this dataset, each sequence in the true positive dataset was shuffled while preserving its dinucleotide distribution. We assume motif occurrences to be homogeneous in length.

### 3.2 Validation on the Rna3Dmotif dataset

#### Validating searches on sequences with known structure

A first aspect to validate is the ability of our method to retrieve the module when the native secondary structure is provided, ensuring the availability of a suitable loop for the module. For this test, the sequences were obtained from the positive dataset, and the structures accommodating their respective modules were generated with RNAfold hard constraint folding. As expected, structure-informed BP2 recovers every existing module.

#### Joint prediction of secondary structure loops and module occurrences

To assess the performance of BayesPairing 2 on sequences of unknown structure, we performed two-fold cross-validation on 100 randomly sampled unique sequences (or on all sequences when fewer were available), for each module, amounting to a total of 6380 sequences. For each sequence-module pair, the candidate with highest score *S* through 20000 sampled structures was considered a **true positive (TP)** if its match score *MS* was above the score cutoff *T* = −2.16, and if its predicted position matched its real three-dimensional structure location. A sequence containing a module on which no accurate prediction was called above the cutoff was considered a **true negative (TN)**. We tested all *λ* values between 0 and 1 and cutoff values between −10 and 10, and found dataset-dependent optimal values of *λ* = 0.35 and a cutoff of −2.16 for this dataset. For the *top 5 scores* sensitivity, any correct prediction within the top 5 candidates could be considered a **TP**, whereas the *top score* test only accepted the highest score output. We also report the F1 score, the Matthews correlation coefficient (MCC), and the false discovery rate (FDR) associated with this cutoff. Formaly the equations of those scores are:

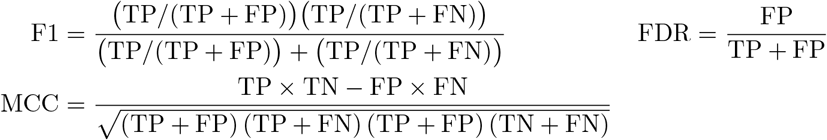

#### Prediction score distribution and false discovery rate

We executed the same two-fold cross-validation experiment on the shuffled sequences described in section 3.1. BayesPairing 2 found no hit on 92% of the 6380 sequences. It should be noted that it is not impossible for a shuffled sequence to contain a good hit for a module.

We obtained distributions of true and false hit scores from the cross-validation dataset. The score distributions, presented in Figure 3a, are clearly distinct, and a score cutoff of −2.16 produced a false discovery rate of 0.061, as reported along with other common metrics in Table 1.

**Table 1:** BayesPairing 2 module identification accuracy on Rna3Dmotif dataset

|  | F1 score | MCC | FDR | Sensitivity<br>(top score) | Sensitivity<br>(top 5 scores) |
| --- | --- | --- | --- | --- | --- |
| **BayesPairing2**<br>**performance** | 0.932 | 0.863 | 0.061 | 0.745 | 0.855 |

**Fig. 3:**
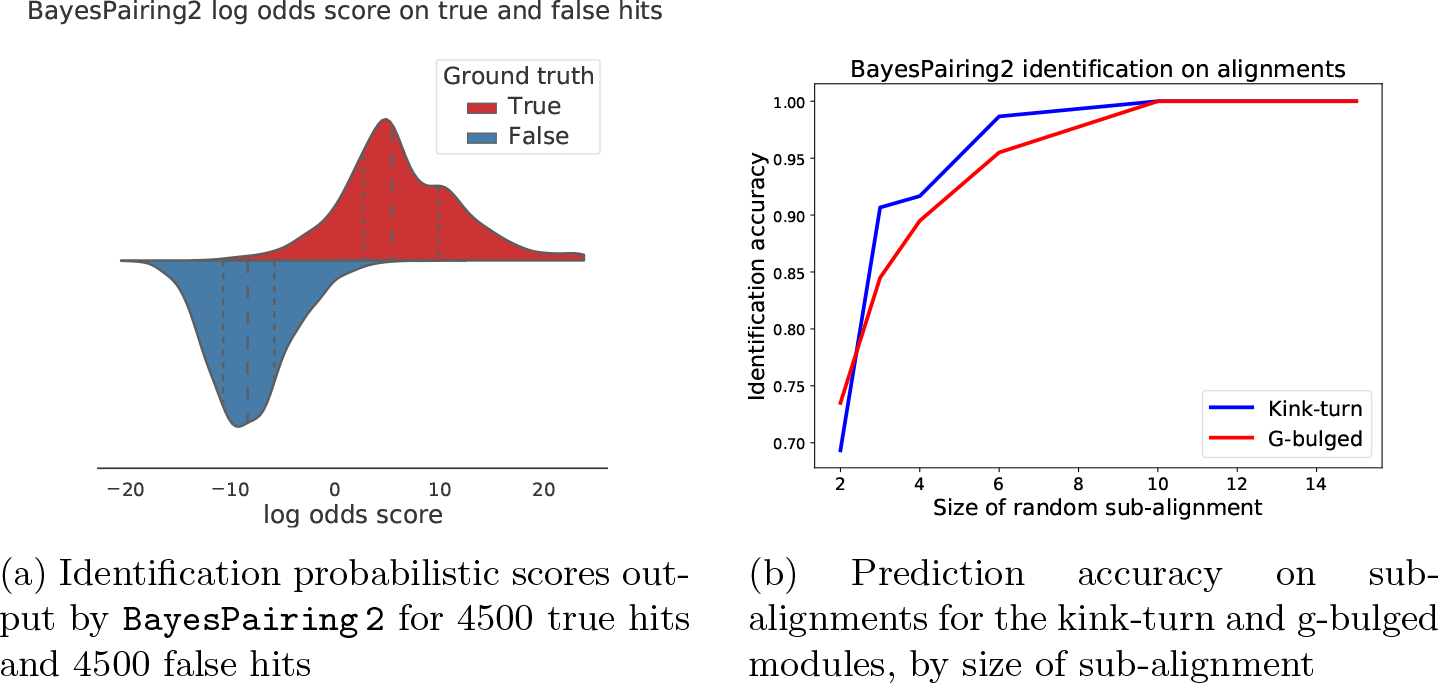
Evaluating BayesPairing 2 scores and accuracy.

### 3.3 Validation on known module alignments from Rfam

#### Sequence search

To complement our cross-validation experiments, we also tested BayesPairing 2 on Rfam alignments of the kink-turn and G-bulged internal loop modules. In these experiments, the modules were associated with their respective families through the Rfam motif database, then trained on one family and tested on the other. The results, for BayesPairing 1 and BayesPairing 2, are displayed in Tables 2a and 2b. We used standard parameters and selected the cutoffs associated to the same false discovery rate of 0.1 for both methods.

**Table 2:**
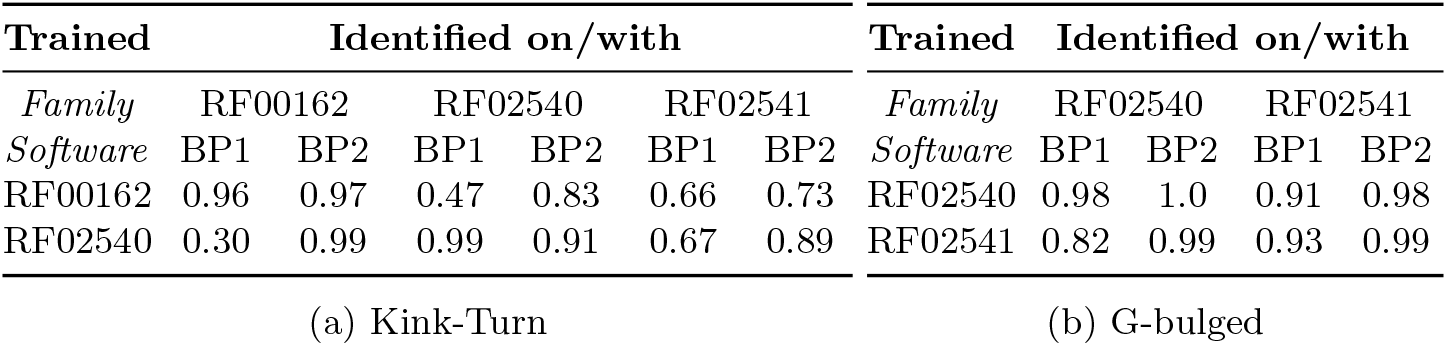
Rfam cross-family results for kink-turn (left) and G-bulged (right)

As observed in section 3.2, BayesPairing 2 is slightly weaker at identifying modules with a strong sequence signal than BayesPairing 1, but considerably stronger when there is significant sequence variation as its signal appears to be more robust. This is particularly well illustrated by the capacity of BayesPairing 2 to identify the ribosomal kink-turn module on SAM riboswitch sequences. While the considerable sequence difference between the ribosome and riboswitch causes a sharp drop of 47% in BayesPairing 1 accuracy when predicting off-family, BayesPairing 2 only loses 25%.

#### Alignment search improvement

Despite positive results in module identification on sequences taken from Rfam, sequence-based methods cannot fully take advantage of the common structure of an alignment. We show the relevance of including module identification on alignments in BayesPairing 2 by improving the results presented in section 3.3. If, instead of parsing individual sequences for modules, we parse randomly sampled sub-alignments, the predictions rise with the size of the sub-alignment until they reach 100%, up from 50 to 95% with sequence predictions by both software tools. Despite very low sample size (500 secondary structure sampled with RNAalifold), the alignment quickly outperforms the sequence predictions for all modules, on all tested families, as shown in Figure 3b.

### 3.4 Time benchmark

The execution time of BayesPairing 2 was measured on 15 sequences (average size of ~ 200 nucleotides) containing a module each, with 5 hairpins, 5 internal loops and 5 multi-branched loops. We searched for 1, 3, 9 and 15 modules, and the execution time as a function of the sequence length and number of modules is displayed in Figure 4. While the software typically requires 2-3 seconds to identify a module in a sequence of length 200, increasing the number of modules searched by a factor of fifteen only doubles its execution time.

**Fig. 4:**
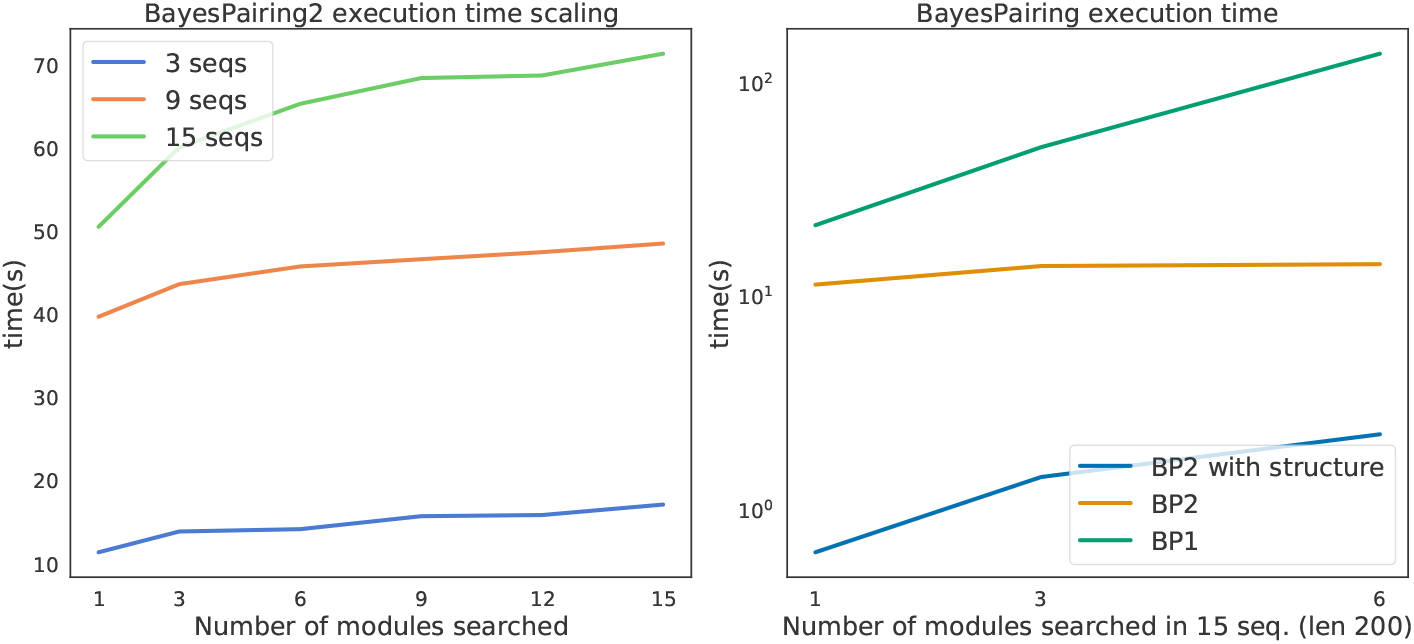
Execution time of BayesPairing 2, as a function of numbers of modules and sequences (left), and compared to BayesPairing 1 (right)

**Fig. 5:**
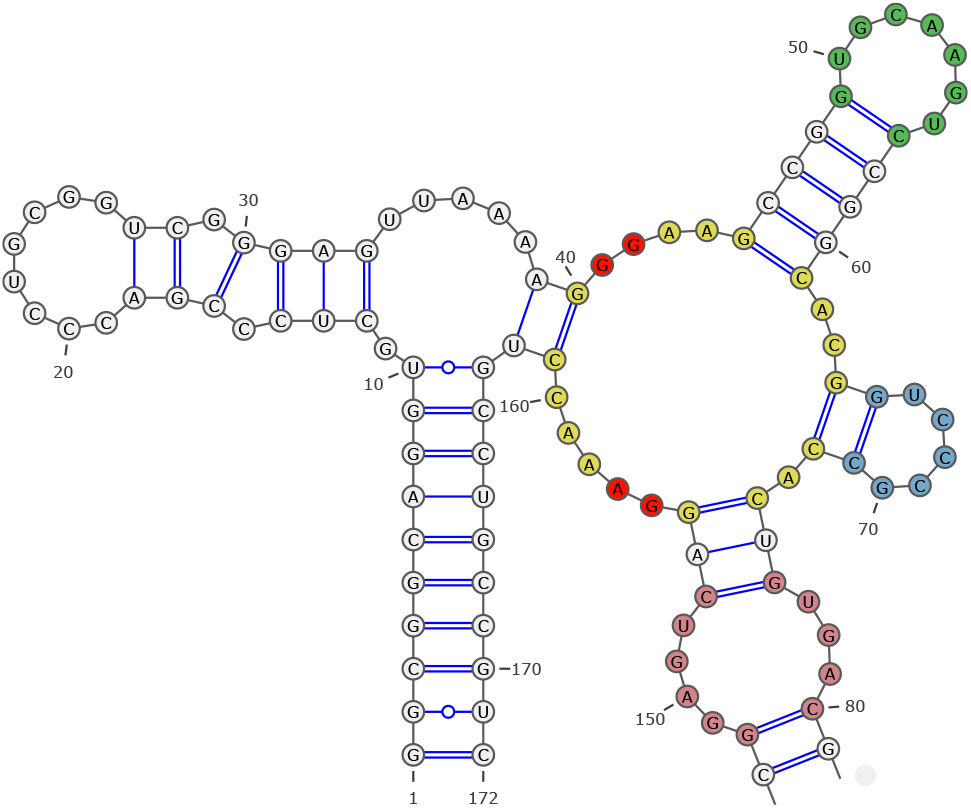
The cobalamin riboswitch four-way junction (in yellow) in PDB structure 4GXY [26]. The adjacent structural motifs used to refine the structural search are highlighted. Bases within 3 Angstrom of the cobalamin molecule in the bound structure are indicated in bright red. Other colors highlight distinct modules

Tests were executed on an Intel(R) Xeon(R) CPU E5-2667 @ 2.90GHz, Ubuntu 16.0.4 with 23 cores, with a total physical memory of 792 gigabytes.

### 3.5 Comparison to the state of the art

The first software to tackle the specific task of identifying 3D motifs in full RNA sequences was RMDetect (2011) [8], which showed good accuracy but was severely limited in the variety of motifs it could identify. BayesPairing 1 improved on this method by adding more flexibility and improving its search efficiency [32]. Another method, JAR3D, does not undertake full sequence searches but scores hairpin and internal loops against a database of models from the RNA 3D Motif Atlas. BayesPairing 2 can be adapted to fulfill the same task, and their purposes are close enough to be comparable. Because BayesPairing 1 has been shown to be a clear improvement on RMDetect, we focus our comparison on the former and JAR3D.

The good performances of BayesPairing 1 [32] relies on the assumption that the structural motif searched has a strong sequence signal. Indeed, the tool identifies motif location candidates through regular expressions. Thus, BayesPairing 1 struggles with motifs trained on a large number of distinct sequences with no dominant sequence pattern.

While it performed well on structure-based datasets with high sequence conservation, our Rfam-based dataset, with an average of 268 sequences from multiple Rfam families for each module, appears challenging for the method and is clearly outperformed by BayesPairing 2 on the dataset described in section 3.1, as shown in Table 3. We also show in Figure 4 that BayesPairing 2 scales much better in the number of modules searched.

**Table 3:** Performances of BayesPairing versions on Rna3Dmotif dataset

|  | F1 score | MCC | FDR | Sensitivity<br>(1 candidate) | Sensitivity<br>(5 candidates) |
| --- | --- | --- | --- | --- | --- |
| BP1 | 0.715 | 0.510 | 0.178 | 0.219 | 0.348 |
| BP2 | **0.932** | **0.863** | **0.061** | **0.745** | **0.855** |

**Table 4:** BayesPairing 2 and JAR3D performances on hairpins (363 seq. in 33 loops), and internal loops (127 seq. in 28 loops) from the RNA 3D Motif Atlas.

| Software | Average Identification TPR and FDR on RNA 3D Motif Atlas |  |  |  |
| --- | --- | --- | --- | --- |
| *Loop type* | Hairpin Loops |  | Internal Loops |  |
| *Software* | TPR | FDR | TPR | FDR |
| **BayesPairing2** | **0.9819** | **0.0020** | **1.00** | **0.0016** |
| **JAR3D** | 0.9685 | 0.0509 | 0.957 | 0.0205 |

JAR3D was also shown to outperform RMDetect in the identification of new variants of RNA 3D modules [40]. However, it does not perform a search on the input sequence, but only takes loops as input. As such, it executes a task that only accounts for a small proportion of BayesPairing 2’s execution time. Indeed, scoring a loop against a model is very rapid, and both tools can score 10,000 module candidates in less than 10 seconds, while the total runtime of BayesPairing 2 when searching for motifs in a single sequence of length 200 is greater than ~ 40 seconds. Therefore, we focus our comparison between BayesPairing 2 and JAR3D on true positive rate and false discovery rate, which contribute to the overall performance of both software.

In order to compare the software, we isolated the scoring component of BayesPairing 2, a function which takes as input a loop and a module and returns a match score between the two, the same input and output as JAR3D. We trained BayesPairing 2 on 51 motifs from the RNA 3D Motif Atlas, including 28 internal loops and 33 hairpin loops. Motifs which constituted full loops and only had occurrences of the same size, the two core assumptions of BayesPairing 2, were selected. Then, internal loops with fewer than three occurrences, and hairpin loops with fewer than 5 occurrences were removed from the dataset. True positive rates (TPR) were computed from predictions on RNA 3D Motif Atlas sequences. False discovery rates (FDR) were estimated from averaged predictions on 100 random sequences per true positive sequences (total 49000). Each random sequence was generated from the nucleotide distribution of the true positive sequences for that module. Default cutoffs were used. For BayesPairing 2, a cutoff of 3.5 was obtained by repeating the process presented in Section 3.2 after setting the weight of the secondary structure to 0, as the secondary structure is only considered in the context of the full sequence which is not part of the input for this specific task. The results are presented in table 4. While the two software present comparable sensitivities, BayesPairing 2 achieves this high sensitivity with higher specificity.

## 4 Discussion

### Applications

The most obvious application for an efficient and parallelizable motif identification framework is to parse sequences for local 3D structure signal. Modular approaches for RNA 3D structure construction like RNA-MoIP [29] have been shown to successfully take advantage of local tertiary structure information. In particular, RNA-MoIP leverages 3D module matches to select the most stable secondary structures to use as a scaffold for the full structure. Indeed, secondary structures that can accommodate known 3D modules are often more predictive of the real structure than those who cannot [8]. To this day, RMDetect, BayesPairing 1 and BayesPairing 2 are the only known full sequence probabilistic module identification tools to be able to identify hairpins, internal loops and junctions, which are key components of many well-known structures, namely several riboswitches. Of the three, BayesPairing 2 is the most scalable. This scalability is essential as many datasets include hundreds of modules [27,30], and this number will keep increasing as more structures are crystallized and mining methods improve.

While the tertiary structure signal encodes information that can be leveraged to build a full 3D structure, its implied functional significance can be taken advantage to refine tasks like sequence classification. Traditional methods for sequence classification include k-mer based techniques [38], as well as sequence and structure motifs [39], but those only use the sequence and secondary structure signals. 3D modules are highly complementary to those methods.

### Identifying multi-branched loops in sequences; applications to riboswitch discovery

One of the distinctive characteristics of BayesPairing 2 is its ability to identify multi-branched loops. These motifs happen to be very common in riboswitches, in which they are often closely related to function, namely in the tyrosine pyrophosphate (TPP) riboswitch, the Cobalamin riboswitch, and the S-adenosyl methionine I (SAM-I) riboswitch [33]. We can use sequences from Rfam riboswitch families to train 3D module models, and then use those models to label new sequences as putative riboswitches.

The software also provides insight on the role of those of 3D modules in the folding dynamics of the riboswitch. Because BayesPairing 2 searches secondary structure ensembles for loops matching known structural modules, it can be used to observe, within the assumptions of the RNAfold library, how easily riboswitch sequences appear to fold into their junction. For instance, the TPP riboswitch’s junction is very present in its Boltzmann ensemble, as its small (13 bases) three-way junction was correcetly identified by our software on 81% of the sequences from the TPP Rfam family.

Because we could hypothesize the frequency of identification of a specific loop to be correlated with its size, it could be expected that the SAM-I riboswitch four-way junction, which counts 28 bases, would be identified less frequently. This is indeed the case as it was identified on 35% of the sequences of its family with a similar pipeline.

The much smaller (17 bases) cobalamin riboswitch junction would then be expected to be found with a frequency somewhere in between 35% and 81%, based on this size assumption. Surprisingly, it was only successfully identified on 3.5% of the Rfam cobalamin family sequences.

However, interestingly, identifying small structural modules (two hairpins and one internal loop) around the junction with a first run of BayesPairing 2 and then using the position of those modules as constraints for a second run raises the frequency of identification of the multi-loop to 32%. The more adjacent motifs are found, the higher the identification confidence was observed to be.

In contrast, applying the same method to the SAM riboswitch, or on shuffled cobalamin riboswitch sequences, does not leave to a significant improvement.

This difference in behavior between riboswitches could be rooted in different factors like co-transcriptional folding, RNA-RNA and RNA-protein interactions and/or the intrinsic difficulty of predicting riboswitch structural element with models learned from bound structures. However, the contrast between the constrained and unconstrained results in the cobalamin riboswitch tends to indicate that some, but not all multi-branched structure are strongly correlated with surrounding loops conformations.

### Limitations and Future Work

Our approaches presents two main limitations. First, the assumption that motif occurrences have a consistent size is not a trivial one to make. For small modules, it is a reasonable assumption that the vast majority of occurrences will have the same size since adding or removing a base would have a large impact on the local 3D structure. However, for larger motifs, and especially junctions, the size constraint can prevent us from identifying some variants. This is something we alleviate in BayesPairing 2 by allowing imperfect matches, with a tolerated difference of up to one base per strand, but further work remains to be done to fully identify motifs bigger than 20 bases, for which this fuzzy matching might not be sufficient.

Second, a consequence of searching secondary structures before sequence is that in the rare cases when the sequence is better conserved than its secondary structure, the accuracy of the tool will suffer. It could however be argued that not overfitting to currently known sequences could be worth losing a bit of accuracy, although this can only be evaluated quantitatively as new structures and module occurrences become available, since the current structure datasets do not show sufficient sequence variability.

Interestingly, a large majority of the modules that cannot be predicted from sequence only by BayesPairing 2 occur in secondary structures that are never generated by RNAsubopt. In many of those cases, a base pair stacking was removed to allow the insertion of the module, at a considerable energy cost. We hypothesize that those small modifications, although not energetically favorable at the secondary structure level, are stabilized by 3D interactions which cannot be inferred from sequence. Going further with this hypothesis, differences in performances are then indicative of the stabilizing effect of non-canonical modules. This assumption could be tested in the future using coarse-grain molecular dynamics to correlate those two metrics.

The other notable limitation of the method is that the loop-based module definition used in our study does not allow the prediction of pseudoknots, nor canonical helices.

## 5 Conclusion

We presented BayesPairing 2, a software for efficient identification of RNA modules in sequences and alignments. BayesPairing 2 strictly outperforms its previous version in execution time, search on provided secondary structures, and sequence search accuracy. It also appears to have complementary strengths to JAR3D, the state of the art for scoring. Finally, its structure-based approach brings a perspective on the place of the motif in the sequence’s Boltzmann ensemble. This added context helps improve identification accuracy, but also the interpretation of the results, and can provide additional information about the role of a module in the folding process. Moreover, the time complexity improvement opens new doors for genome-wide sequence mining for local 3D structure patterns. As new RNA structures and sequences become available, more modules will be discovered, and BayesPairing 2 is fast enough to take advantage of its customizability to contribute to filling the gap between secondary and tertiary structure prediction tool by associating a wide selection of RNA modules of interest to those new sequences.

## Acknowledgements

The authors are greatly indebted to Anton Petrov for providing us with alignments between RNA PDB structures and Rfam families, which helped us match 3D modules to sequence alignments.

